# Coiled-coil-mediated phase separation of Spef1 for non-centrosomal microtubule organization and function

**DOI:** 10.1101/2025.07.31.667833

**Authors:** Jinqi Ren, Juyuan Liu, Dong Li, Xueliang Zhu, Wei Feng

## Abstract

Central-pair microtubules (CP-MTs) are non-centrosomal MTs essential for planar beat pattern of cilia. The CP-MT formation requires the MT-associated protein Spef1, but the underlying molecular mechanism remains unclear. Here, we show that Spef1 undergoes liquid-liquid phase separation (LLPS) to facilitate non-centrosomal MT assembly by enriching tubulins. The LLPS of Spef1 is mediated by its C-terminal coiled-coil (CC) domain. Crystallography reveals that the Spef1-CC domain forms a parallel CC dimer with a unique charge distribution pattern on the surface. The dimerization capacity and charge distribution of Spef1-CC are both critical for controlling *in-vitro* LLPS. Disruption of the dimerization capacity abolishes ciliary functions of Spef1. In contrast, a charge-changing mutant with attenuated LLPS still supports the CP-MT formation but results in cilia with abnormal beat pattern. Thus, the CC-mediated LLPS of Spef1 provides a mechanistic explanation for its prominent role in controlling non-centrosomal CP-MT organization and function in the axoneme.

**Significance statement:** The MT-associated protein Spef1 is a new essential player for the non-centrosomal CP-MT formation in motile cilia and flagella. This study reveals the unexpected LLPS feature of Spef1, leading to forming biomolecular condensates that enrich tubulins to facilitate non-centrosomal MT assembly. Spef1-CC contains a unique charge-distribution pattern, together with its dimerization capacity, contributing to multivalent interactions for initiating LLPS. The LLPS property of Spef1 is important for CP-MT formation and Spef1-mediated ciliary function. The formation of Spef1-LLPS condensates indicates that they work as MT nucleation centers and tubulin sources for the continuous growth of CP-MTs or repairing CP-MTs damaged during ciliary beating, and suggests that LLPS may be a common process for generating and organizing non-centrosomal CP-MTs in the axoneme.

## Introduction

Motile cilia and flagella are hair-like organelles featured with an exquisite microtubule (MT)-based structure known as the axoneme ^1, 2^. Through oscillatory beating of the axoneme, motile cilia and flagella can drive cell movement or extracellular fluid flow ^3, 4^. The core MT-scaffold of the axoneme is comprised by nine peripheral doublet MTs that originate from the basal body, a specialized centriole, and a central pair of singlet MTs (CP-MTs) ^2, 5^. In contrast to the doublet MTs, CP-MTs are non-centrosomal MTs and sit above the transition zone without any connection to the basal body ^6^. The two CP-MTs are associated with different proteinous projections to form the CP apparatus. The precise coordination between the CP apparatus and the radial spokes protruding from peripheral doublet MTs allows motile cilia or flagella to beat in a specific pattern ^7, 8^. For instance, mammalian cilia containing CP-MTs beat in a back-and-forth (planar) manner, whereas those lacking CP-MTs beat in a rotatory pattern ^9, 10^. Accordingly, the lack or partial defect of the core MT-scaffold in the axoneme leads to abnormal ciliary beat patterns and causes a group of cilia-related diseases, namely primary ciliary dyskinesia (PCD) in humans ^11, 12^.

CP-MTs are primarily initiated from preassembled MT-seeds or undergo *de novo* self-assembly from enriched tubulins in the axonemal lumen ^6, 13^. The MT-seeds and tubulins for CP-MT formation are either imported by intraflagellar transport or generated by katanin-mediated severing of axonemal MTs ^14–16^. In addition to katanin, Camsap family proteins and the WD40-repeat-containing protein Wdr47 are involved in controlling proper CP-MT formation ^9, 17^. Camsap proteins are katanin-binding partners and can recognize and stabilize MT minus-ends ^18–20^, while the N-terminal domain of Wdr47 specifically recognizes a unique basic-helical motif in Camsap proteins to form a Wdr47-Camsap-katanin axis for assembling CP-MTs ^9, 21^. In this axis, katanin produces preassembled MT-seeds by severing axonemal MTs, Camsap proteins recognize and stabilize these MT-seeds to promote their extensions, and Wdr47 recruits Camsap proteins into the axonemal lumen to ensure the proper number and localization of CP-MTs ^9, 21^. As Camsap proteins mainly stabilize the minus ends, the newly generated CP-MTs are expected to require additional MT-regulators to stabilize and organize due to the inherent dynamic properties of MTs.

Spef1, also called CaLponin-homology And Microtubule-associated Protein (CLAMP), was originally identified as a conserved MT-associated protein in mouse sperm flagella and the inner ear organ of Corti ^22, 23^. Spef1 contains an N-terminal calponin-homology (CH) domain that associates with MTs and a C-terminal coiled-coil (CC) domain responsible for dimerization ^10, 24^. Combining its MT-binding and dimerization capacities, Spef1 tends to function as a cross-linker to bundle MTs and contribute to MT stability ^10, 23, 24^. As a prominent MT-regulator, Spef1 exhibits the capacity of controlling MT-organization in epithelia for planar cell polarity establishment and directional cell migration ^25, 26^. More intriguingly, Spef1 also localizes to the CP and is an essential player for the CP-MT formation in motile cilia. Depletion of Spef1 in mouse ependymal cells (mEPCs) by RNAi leads to the complete loss of CP-MTs in multicilia, resulting in an abnormal rotatory beat pattern ^10^. Mutations in the CH or CC domain to disrupt the MT-binding or dimerization capacity of Spef1 severely impair its ability to restore the CP-MT formation and planar ciliary beating in endogenous Spef1-depleted mEPCs ^10^. Thus, Spef1 is proposed to bundle and stabilize CP-MTs, although the underlying mechanism remains largely unclear.

Interestingly, Spef1 is enriched at the tip of growing cilia and tends to form puncta distributed along CP-MTs ^10, 27^, resembling droplets formed by liquid-liquid phase separation (LLPS). In this study, we characterize Sepf1 and find that it undergoes LLPS to form biomolecular condensates critical for ciliary CP-MT formation and function.

## Results

### Spef1 undergoes LLPS

To investigate the mechanism underlying Spef1-mediated organization of CP-MTs, we initiated this work with the biochemical characterization of full-length Spef1. Gel filtration analysis revealed that the protein exhibited significant heterogeneity under low-salt conditions (150 mM NaCl), as evidenced by the presence of multiple elution peaks (**Figure S1A**). In contrast, under high-salt conditions, the protein eluted as two well-defined peaks (**Figure S1A**), indicating a shift toward a more homogeneous state. This behavior is likely attributable to weak, non-specific electrostatic interactions or transient oligomer formation that are promoted at low ionic strength and effectively suppressed upon salt addition. In addition, the solution of Spef1 becomes turbid with the increase of temperature even at high salt concentrations (**Figure S1B**). In light of current knowledge on protein LLPS, this salt-dependent dissociation phenomenon suggests that Spef1 may transition between distinct solution states to facilitate liquid–liquid phase separation. To evaluate the LLPS capacity of Spef1, we fused an EGFP-tag to Spef1 for the *in vitro* fluorescence-based LLPS assay. During protein sample preparation, the buffer containing a high concentration of salt (500 mM NaCl) was used to prevent the precipitation of Spef1. Upon dilution into the buffer without salt, EGFP-Spef1 (with an estimated concentration of 20 µM) immediately forms oil-like droplets, a characteristic feature of LLPS, at 25°C and the number and size of these droplets increases over time (**Figure 1A-C**). Importantly, fusion events are readily observed (**Figure 1D**), confirming the liquid nature of the droplets. We further performed fluorescence recovery after photobleaching (FRAP) assays and observed rapid recoveries of the fluorescence of droplets after photobleaching (**Figure 1E**), indicating dynamic rearrangement within the droplets and the frequent exchange between the condense and dilute phases, another canonical feature of LLPS. Interestingly, the recovery markedly decreases when Spef1 droplets formed for 4 hours were photobleached (**Figure 1E**), indicating their gradual transit from a liquid to a gel-like state. Moreover, increasing the temperature to 37°C apparently accelerates this process (**Figure 1E**).

**Figure 1:**
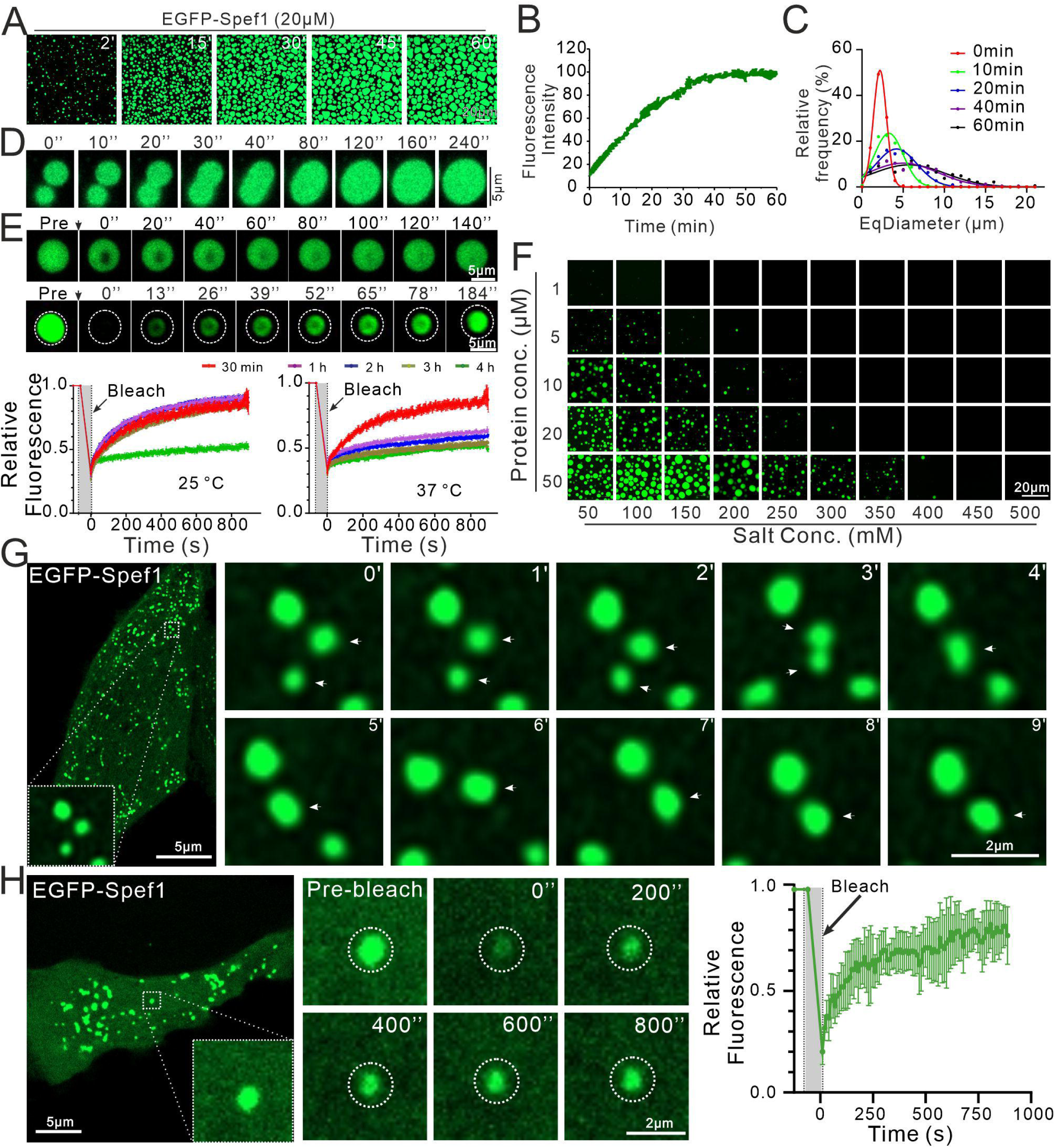
Spef1 undergoes LLPS both *in vitro* and *in cellulo*. **(A)** Representative fluorescence images of EGFP-Spef1 phase separation at indicated time points in buffer with a physiological salt concentration (150 mM NaCl). Protein concentration of EGFP-Spef1 is 20 μM. Scale bar: 20 µm. **(B)** Overall fluorescence intensities of EGFP-Spef1 droplets formed over the time course of 60 min. The maximal fluorescence intensity was normalized to 100% and the values shown are mean ± SD (n = 3 images). **(C)** Equivalent diameter (EqDiameter) frequency distribution of EGFP-Spef1 droplets formed at the indicated time points. **(D)** Fusion of adjacent EGFP-Spef1 droplets with the increase of time. **(E)** FRAP analysis of EGFP-Spef1 droplets. Time-lapse images from the full- and partial-FRAP experiments (upper panel). Fluorescence recovery curve after photobleaching (lower panel). Photobleaching was performed at the indicated time points (red: 30 min; purple: 1 h; blue: 2 h; brown: 3 h; green: 4 h), and the fluorescence recovery was measured at 25°C and 37°C, respectively. Data are presented as mean ± SD. **(F)** LLPS diagram showing the phase separation of EGFP-Spef1 at different protein and salt concentrations. **(G)** Representative fluorescence images of HeLa cells transfected with EGFP-Spef1. Enlarged views in real time of the dashed box in the right panel showing fusion of droplets. White arrowheads indicate the syncretic droplets before and after fusion. **(H)** FRAP analysis of EGFP-Spef1 droplets in HeLa cells. Time-lapse images from the FRAP experiment (left panel). Fluorescence recovery curve after photobleaching (right panel). Data are presented as mean ± SD.

We next generated the phase separation diagrams by gradually changing concentrations of EGFP-Spef1 and salt (**Figure 1F**). At the physiological salt condition (150 mM NaCl), the critical concentration required for the LLPS of EGFP-Spef1 is ∼5 µM (**Figure 1F**). To eliminate potential interferences of the EGFP-tag, we removed it and labeled full-length Spef1 with the fluorochrome FITC. The FITC-labeled and non-labeled Spef1 proteins were then mixed at a molar ratio of 1:10. As expected, tag-free Spef1 forms droplets capable of fusion and rapid fluorescence recovery when entire droplets were photobleached (**Figure S1C-D**), indicating frequent exchange of Spef1 molecules between the droplets and dilute phases. Phase separation diagrams indicate comparable LLPS ability to EGFP-Spef1 (**Figure 1F** vs. **Figure S1E**), indicating that the influence of the EGFP-tag on the LLPS, if any, is negligible. Furthermore, the critical concentration is as low as 0.1 µM in 50 mM NaCl (**Figure S1E**). Taken together, Spef1 is capable of undergoing LLPS to form liquid droplets.

### Spef1 forms biomolecular condensates in cells

Based on the observation from the *in vitro* LLPS assay, we expressed EGFP-Spef1 in different cell lines by transient transfection to assess whether Spef1 can undergo LLPS under cellular conditions. Consistent with previous publications ^10^, exogenous Spef1 localizes to MTs and also formed puncta (**Figure 1G** and **Figure S2**). EGFP-Spef1 appears to distribute along MTs, inducing MT bundling due to its cross-linking activity (**Figure S2**) ^10^, whereas its punctate distributions are also prominent in all cells (**Figure 1G** and **Figure S2**). Living-cell imaging showed that these puncta, whose diameters ranged from 0.2 μm to 0.5 μm in HeLa cells, are capable of rapid fusion (**Figure 1G**). FRAP assays further demonstrated that ∼80% of the fluorescence intensity of the puncta can be recovered after photobleaching with the t_1/2_ of ∼10 s (**Figure 1H**). Thus, Spef1 exhibits the LLPS capacity to form dynamic biomolecular condensates in cells.

### Spef1 condensates enrich tubulins to promote MT assembly

The co-localization of EGFP-Spef1 condensates with MTs in cells indicated that Spef1 could somewhat control the MT-network organization in addition to the MT-bundling. Moreover, previous studies demonstrated that endogenous Spef1 is preferably distributed at the cilia tip in growing cilia ^10, 24^, suggesting its potential role in MT growth. We thus speculated that the newly-discovered Spef1-LLPS condensates might promote MT growth by enriching tubulins for polymerization, although the well-known MT-bundling activity of Spef1 would be also essential. To check the possibility of tubulin enrichment by Spef1-LLPS condensates, we mixed EGFP-Spef1 and TAMRA-labeled tubulins together and performed the LLPS assay by diluting them into the buffer with a low concentration of salt (100 mM KCl). In contrast to the TAMRA-labeled BSA, tubulins co-condense with EGFP-Spef1 to form similar droplets with periphery-enriched distributions (**Figure 2A**), demonstrating the tubulin-enrichment capacity of Spef1-LLPS condensates. To rule out possible artifacts caused by the fluorescent dye, this tubulin-enrichment capacity was further evaluated using a sedimentation assay using both tag-free Spef1 and unlabeled tubulins. As expected, unlabeled tubulins co-sediment with Spef1-LLPS droplets in a concentration-dependent manner (**Figure 2B**). Therefore, Spef1-LLPS condensates are able to enrich tubulins.

**Figure 2:**
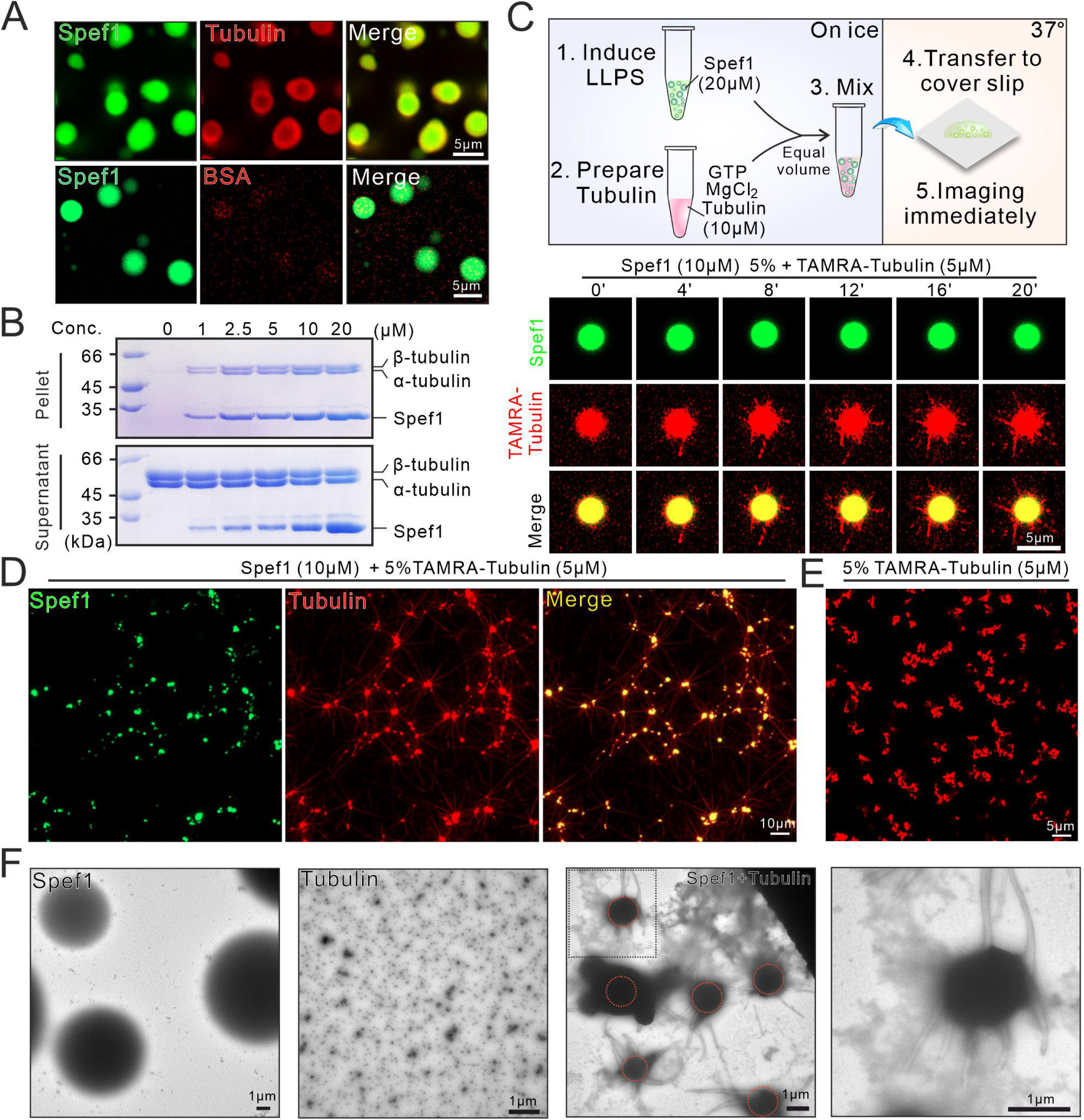
Spef1 condensates enrich tubulins to promote MT assembly. **(A)** Fluorescent images of EGFP-Spef1 (green) condensates prepared with TAMRA-labeled α/β-tubulins or bovine albumin (BSA) (red). TAMRA-labeled BSA was used as a negative control. Scale bar: 5 µm. **(B)** The sedimentation assay in the presence of increasing concentrations of Spef1 and tubulins. **(C)** Time-lapse fluorescent images of Spef1-tubulin co-condensates (green and red, 10 µM and 5 µM, respectively) prepared in the MT-polymerization buffer. Representative of three experimental replicates. **(D-E)** Representative fluorescence micrographs of EGFP-Spef1 (green) and TAMRA-labeled tubulins (red) taken 20 min after starting the reaction (D). For the control experiment, TAMRA-labeled tubulins alone were in the BRB80 buffer (E). **(F)** Negative-stain electron micrographs. The samples were prepared in the MT-polymerization buffer. They were fixed and negatively stained 10 min after starting the reaction. Spef1 and tubulins alone were used as controls. Scale bar: 1 µm.

To investigate whether the high local concentration of tubulins in these droplets would facilitate the nucleation and assembly of MTs, we initially employed the turbidity-based MT polymerization assay to test this hypothesis. However, both condensate formation and MT polymerization resulted in absorbance at 350 nm, making it difficult to attribute the absorbance to MT growth. Then we used time-lapse fluorescence microscopy to monitor the droplets formed by the LLPS of EGFP-spef1 and tubulins (contains 5% TAMRA-tubulin) in the BRB80 buffer (**Figure 2C-E**). The final concentration of tubulins we used is 5 μM, which is below the critical concentration for MT self-nucleation *in vitro* ^28^. With the addition of GTP (an essential factor for MT growth), the droplets are observed to emanate radial MT arrays, or MT-asters, over time (**Figure 2C**). Moreover, these MT-asters tends to further inter-connect to assemble and organize a complex MT-network following the time (**Figure 2D**), indicating an intrinsic capacity of Spef1-LLPS condensates for organizing MTs. Similar results were obtained using unlabeled tubulins stained with the SiR-tubulin (**Figure S3A-B** and **Video S1**), a MT-specific dye ^29^. To assess the properties of these MT-asters, we next utilized negative-stain electron microscopy (EM) to analyze them in detail. As controls, Spef1-LLPS condensates display a spherical structure with a smooth surface, whereas tubulins with concentrations below that for self-nucleation shows no obvious MT formation (**Figure 2F**). Upon mixing Spef1 and tubulins for a short period of time, Spef1-LLPS condensates were found to contain radial MT-bundles (**Figure 2F**), similar to the MT-aster structures under fluorescence microscopy (**Figure 2C**). Thus, the MT-bundling activity of Spef1 and the tubulin enrichment capacity of Spef1-LLPS condensates likely work together to control the thick MT-bundle formation.

### The CC domain of Spef1 controls its phase separation

Based on the knowledge of biomolecular condensates formed by LLPS ^30, 31^, intrinsically disordered regions (IDRs) and CC domains are major determinants for LLPS by mediating multivalent interactions. Spef1, besides the N-terminal CH domain, contains a C-terminal CC domain and an IDR between the two domains (**Figure 3A**). To determine which region in Spef1 is responsible for its LLPS, we generated a series of truncated fragments and assessed their abilities for LLPS. Deletion of the CH domain has little impact on the droplet formation, while deletion of IDR greatly decreases protein solubility, as evidenced by the high aggregation of EGFP-Spef1-ΔIDR (**Figure 3B-C**). In contrast, the removal of the CC domain completely abolishes the ability of LLPS (**Figure 3B-C**), signifying the crucial role of the CC domain for initiating LLPS. Consistently, neither EGFP-CH nor EGFP-IDR drives LLPS, whereas the FITC-labeled CC domain alone undergoes LLPS under the same conditions (**Figure 3B-C**). To assess whether the CC domain phase separates through dimerization, we replaced this domain with the CC domain of GCN4 (GCN4-CC), a canonical dimerization domain. Nevertheless, C-GCN4, the Spef1 chimera containing GCN4-CC, fails to show LLPS (**Figure 3B-C**), suggesting that the CC domain of Spef1 (referred to as Spef1-CC) does not phase separate solely through its dimerization ability.

**Figure 3:**
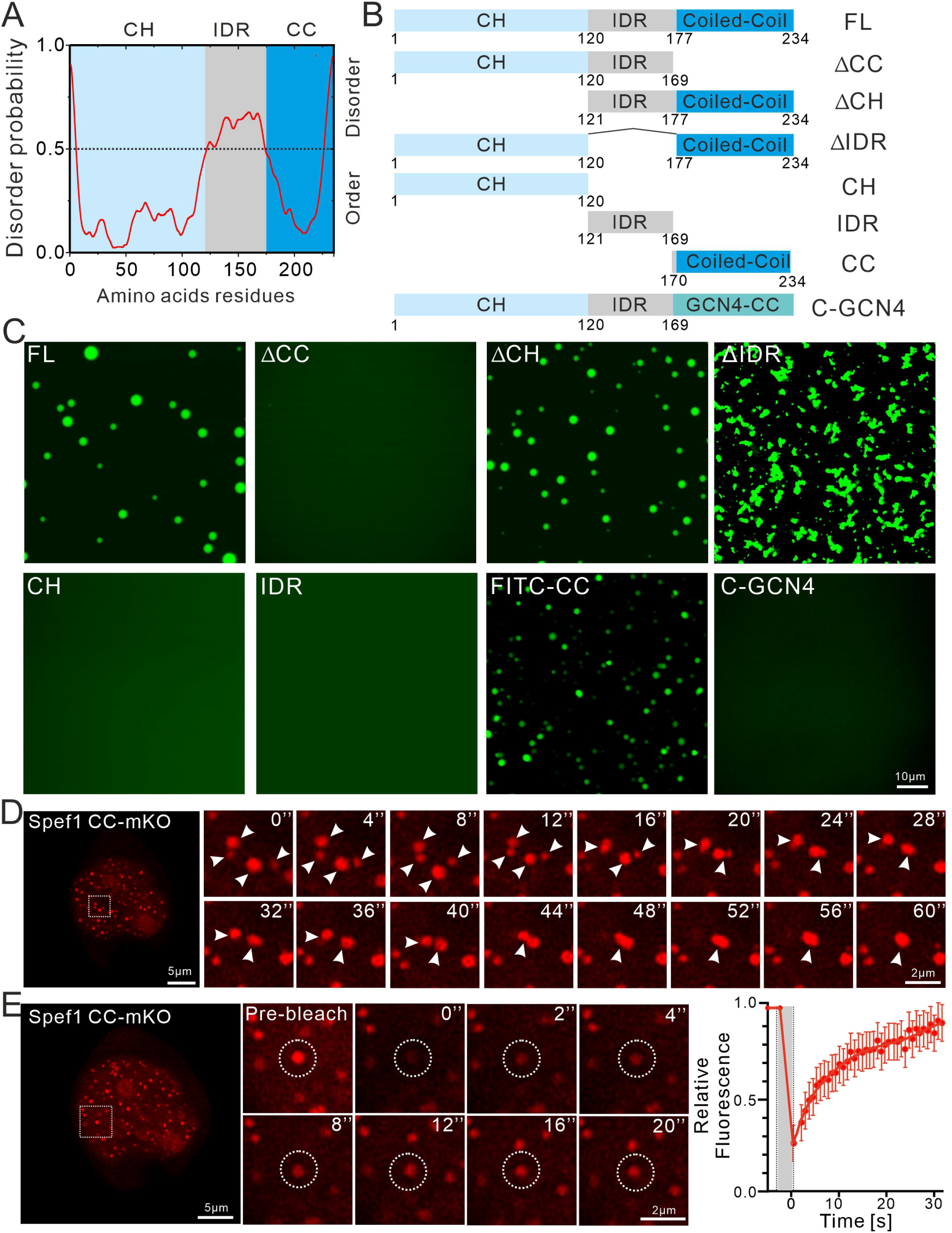
The CC domain of Spef1 controls its phase separation. **(A)** Schematic representation of the domain organization (top) and sequence feature (bottom) of Spef1. CH, Calponin homology domain; IDR, intrinsic disordered region; CC, coiled coil domain. The line at 0.5 (y axis) is the cut-off for the disorder (>0.5) and order (<0.5) predictions. **(B)** Schematic diagram of the truncation mutants and chimeras of Spef1 used in this study. **(C)** *In vitro* phase separation behaviors of the truncation mutants and chimeras of Spef1. Representative fluorescence images of the LLPS ability of the different truncation mutants and chimeras of Spef1. Scale bar: 10 µm. **(D)** Representative fluorescence images of HeLa cells transfected with Spef1-CC containing a C-terminal fluorescent marker (mKO). Enlarged views in real time of the dashed box in the right panel showing fusion of droplets. White arrowheads indicate the syncretic droplets before and after fusion. Scale bar: 5 µm. **(E)** FRAP analysis of Spef1-CC-mKO droplets in HeLa cells. Time-lapse images from the FRAP experiment (left panel). Fluorescence recovery curve after photobleaching (right lanel). Data are presented as mean ± SD. Scale bar: 5 µm.

We next verified whether Spef1-CC is able to undergo LLPS in cells by expressing it tagged at the C-terminus with the fluorescent protein monomeric Kusabira Orange (mKO) in HeLa cells. Spef1-CC-mKO obviously forms a large number of puncta capable of frequent fusion and rapid recovery after photobleaching, though it tends to accumulate in the nucleus (**Figure 3D-E**). Taken together, Spef1-CC is the critical region sufficient for driving LLPS *in vitro* and *in cellulo*.

### Structure of Spef1-CC

Given that Spef1-CC is the LLPS-inducing region that may possess special characteristics, we next attempted to biochemically and structurally characterize this domain to reveal the mechanism for driving LLPS. Although Spef1-CC easily undergoes LLPS under general buffer conditions, the high concentration of salts stabilized the protein in a homogeneous state (**Figure S4A**). Utilizing a high-salt buffer, we conducted the crystal screening for this domain and eventually obtained the high-quality crystals suitable for structural determination after extensive trials. The structure of Spef1-CC was determined by the molecular replacement method and refined to 2.3 Å (**Figure 4A** and **Table S1**). A total of six molecules were found in the asymmetric unit to form three parallel CC dimers that could be well superimposed with little difference (**Figure S4B-C**), demonstrating that Spef1-CC adopts a canonical dimeric conformation. The Spef1-CC dimer consists of seven heptad repeats (*a*-*g*) and spans with a length of ∼8 nm (**Figure 4A-B**). The two helices in the Spef1-CC dimer are held together by the hydrophobic packing formed by the hydrophobic residues from *a* and *d* positions (i.e., I184, L191, V198, L201, V205, L208, L212, I219, L222 and L226) (**Figure 4B-C**). More intriguingly, the electrostatic surface analysis of the Spef1-CC dimer revealed that it possesses a distinctive charge distribution pattern on the surface (**Figure 4D**). From one side of the Spef1-CC dimer, the N-terminal halves of both helices are enriched with a negative charge potential contributed by E186, E188, E190 and E196 (**Figure 4C** and **Figure S4D**). In contrast, the C-terminal half of one helix is enriched with a positive charge potential contributed by K204, R207, K215 and R218, while that of the other helix is neutralized with the mixture of the positively and negatively charged residues (**Figure 4C** and **Figure S4E**). The other side of the Spef1-CC dimer (with a rotation of ∼180°) contains the similar charge distribution pattern with the unique enrichment of the positive and negative charge potentials (**Figure 4D**). This electrostatic surface analysis suggests that the electrostatic interactions would happen between the two halves of the Spef1-CC dimer due to their complementary charge potentials (**Figure 4D-E**). Thus, the unique charge distribution pattern of the Spef1-CC dimer would provide a further layer of inter-molecular interactions, in addition to dimerization, to facilitate the multivalent interactions for inducing LLPS (**Figure 4E**). Primary sequence analysis reveals that the CC domain is highly conserved in Spef1 across different species from protists to mammals (**Figure 4C**), suggesting that the CC-mediated LLPS may be a common feature of Spef1 orthologues.

**Figure 4:**
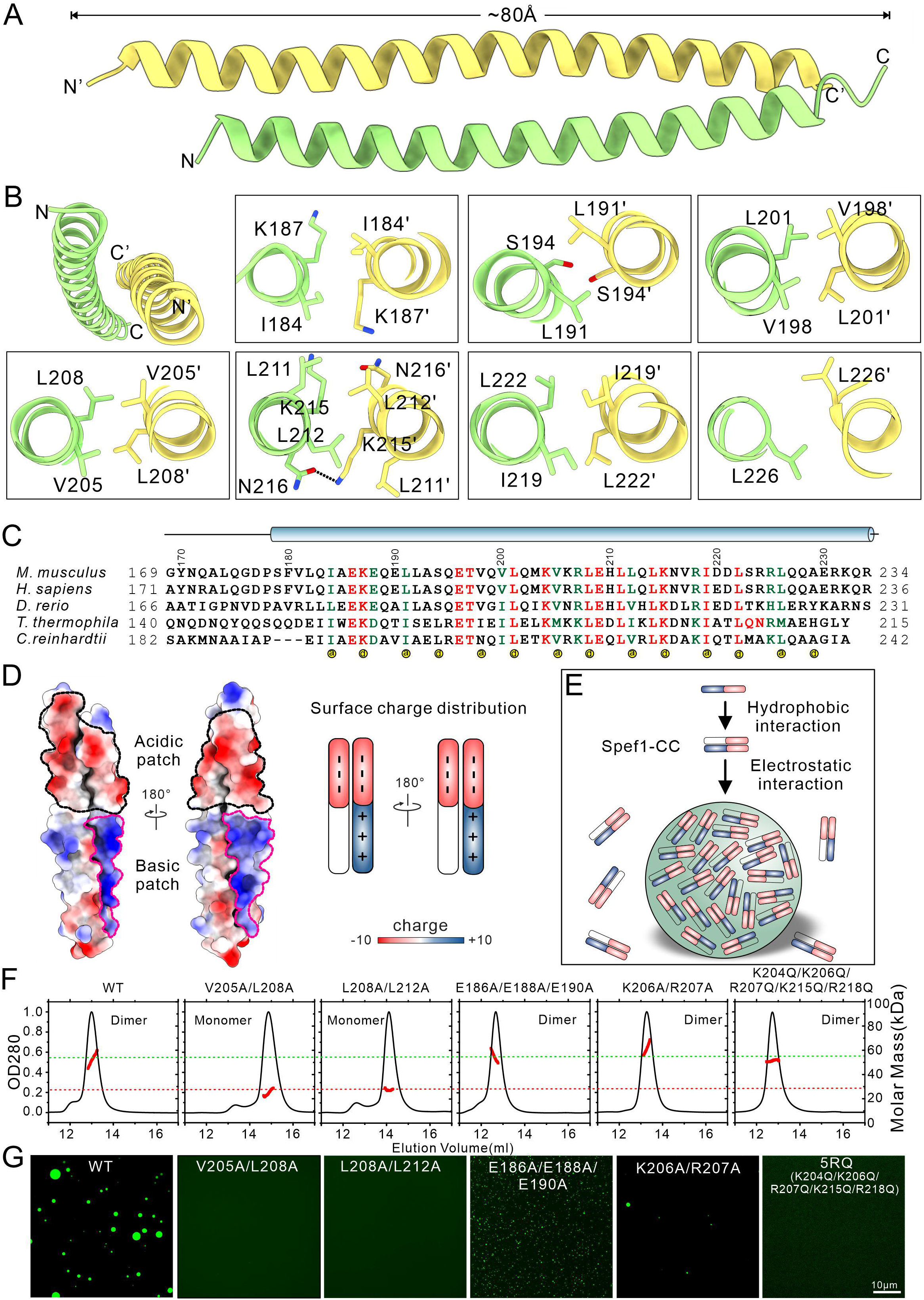
Structure of Spef1-CC and its intrinsic special properties for phase separation. **(A)** Overall structure of Spef1-CC dimer. In this drawing, one chain is colored in yellow and the other is colored in green. Two monomers interact with each other in a parallel fashion. **(B)** Structural details of the dimerization interface for the Spef1-CC dimer. Hydrogen bonds are indicated by dashed lines. **(C)** Sequence alignment of Spef1-CC from different species. The NCBI accession numbers of these proteins involved are as below: mouse (*M. musculus*, NCBI Reference Sequence: NP_081917.1), human (*H. sapiens*, NCBI Reference Sequence: NP_056232.2), fish (*D. rerio*, NCBI Reference Sequence: NP_956821.1), green alga (*C. reinhardtii*, NCBI Reference Sequence: XP_001695399.2) and Tetrahymena (*T. thermophila*, NCBI Reference Sequence: XP_001009857.2). Identical residues and highly conserved residues are colored in red and green, respectively. The essential residues responsible for the hydrophobic contacts in the inter-domain interfaces are marked with yellow dots at the bottom. **(D)** Electrostatic surface potential of the Spef1-CC dimer showing the special charge distribution pattern. The protein surface is colored according to its electrostatic potential from red (negatively charged) to blue (positively charged). **(E)** A schematic model showing the weak electrostatic interaction mediated-LLPS of Spef1-CC. **(F)** Biochemical characterization of the oligomeric states of wild-type Spef1 and its mutants by SEC-MALS. The calculated molecular weight of wild-type Spef1 matches a dimeric state. **(G)** *In-vitro* phase separation behaviors of different Spef1 mutants with mutations in the CC domain to dissociate the dimer or disrupt the charge distribution. The protein concentration used for inducing LLPS is 10 μM. Scale bar: 5 µm.

### Intrinsic special properties of Spef1-CC drive phase separation

To evaluate whether the intrinsic properties of the Spef1-CC dimer are essential for inducing LLPS, we created point mutations either in the CC packing interface to disrupt the dimerization or on the surface to change the complementary charge potentials of Spef1-CC in full-length protein (**Figure 4B-D**). We assessed the oligomeric states of these mutants in the buffer with the high concentration of salt by the size-exclusion chromatography coupled with multi-angle light-scattering (SEC-MALS) assay. As expected, in comparison to the dimeric state of the wild-type protein, the CC-disrupting V205A/L208A and L208A/L212A mutants adopt a monomeric state in solution, while the charge-changing E186A/E188A/E190A and K206A/R207A mutants still exist as a dimer due to the little impact of the mutations on the surface for CC dimer formation (**Figure 4F**). In the fluorescence-based LLPS assay, the V205A/L208A and L208A/L212A mutants show no capacity of LLPS to form droplets (**Figure 4G**), indicating that these CC-disrupting mutations abolish the LLPS ability of Spef1. On the other hand, the E186A/E188A/E190A and K206A/R207A mutants still form droplets albeit with a much smaller size in comparison to the wild-type protein (**Figure 4G**). Thus, we created a more dramatic charge-changing mutant that contains five point mutations (K204Q/K206Q/R207Q/R215Q/R218Q, 5RQ in short) on the surface (**Figure 4C-D**). The 5RQ mutant also exists as a dimer in solution but exhibites a reduced capacity of LLPS compared to the other two charge-changing mutants (**Figure 4F-G**). However, these five mutations do not completely abolish its ability to undergo LLPS, as the 5RQ mutant can still form droplets at high protein concentrations (**Figure S5A**). Given the more severe impairment induced by the CC-disrupting mutations (2-point *vs.* 5-point), the CC-mediated dimerization seems to a prerequisite for LLPS and the surface charge distribution is most likely to be an auxiliary factor to further promote LLPS. We also evaluated the MT-bundling abilities of these mutants by *in vitro* MT-bundling assays. Consistent with previous results, the wild-type Spef1 induced massive MT bundling at the concentration of 4 μM. By contrast, both the 5RQ and L208A/L212A mutants failed to effectively bundle MTs at the same concentration (**Figure S5B**), suggesting a potential correlation between LLPS and the MT-bundling ability of Spef1.

To corroborate the *in-vitro* LLPS results of different Spef1 mutants, we expressed the 5RQ and L208A/L212A mutants in HeLa cells. Consistently, none of them forms obvious biomolecular condensates as the wild-type Spef1 does (**Figure S5C-D** *vs.* **Figure 1G-H**), indicating that these point mutations also severely impact the formation of Spef1-LLPS condensates in cells. Importantly, this effect is not due to instability of the mutant proteins, as western blot analysis shows that both mutants are expressed at levels comparable to the wild-type Spef1 (**Figure S5E)**, indicating similar protein stability. Moreover, since the Spef1-LLPS condensates can enrich tubulins and promote MT growth (**Figure 2**), we incubated these Spef1 mutants with tubulins and checked the MT formation. As expected, no MT-aster structures were observed (**Figure S5F-G**), demonstrating that the Spef1 mutants with the mutations in Spef1-CC lose the capacity of controlling MT organization. Taken together, the intrinsic special properties of Spef1-CC are essential for driving LLPS of Spef1 and the related function in the control of MT growth and assembly, and the dimerization capacity and charge distribution pattern of Spef1-CC are both phase-separation determinants for multivalent interactions of Spef1 to initiate LLPS.

### Ciliary function of Spef1 requires its LLPS property

We next wondered whether the LLPS of Spef1 also plays a role in the CP-MT formation. We depleted Spef1 in cultured mEPCs using a previously validated siRNA (Spef1i) ^10^ and performed rescue experiments through lentivirus-mediated ectopic expression of GFP-tagged wild-type Spef1 (WT) or its 5RQ, K206A/R207A (KR) and L208A/L212A (LL) mutants (**Figure 5A-B**). Consistent with previous report ^10^, live imaging indicated that, while cilia in 93.6% of mEPCs treated with a negative control siRNA (NCi) mainly displayed a planar beat pattern, those in 79.8% of mEPCs treated with Spef1i became rotatory (**Figure 5C-D**). Furthermore, cilia in 90.2% of GFP-WT-positive mEPCs treated with Spef1i beat planarly (**Figure 5C-D**), indicating a full rescue of the defect. In contrast to the WT, cilia in 75.4% of GFP-KR-positive and 45.6% of GFP-5RQ-positive mEPCs displayed planar beat, whereas the expression of GFP-LL did not significantly alter the beat patterns of Spef1i-treated mEPCs (21.8% vs. 20.2% for cells with planar ciliary beat) (**Figure 5C-D**). Therefore, the KR and 5RQ mutants display progressively decreasing abilities to restore the planar beat pattern of Spef1i-treated cilia, whereas the LL mutant is inactive.

**Figure 5:**
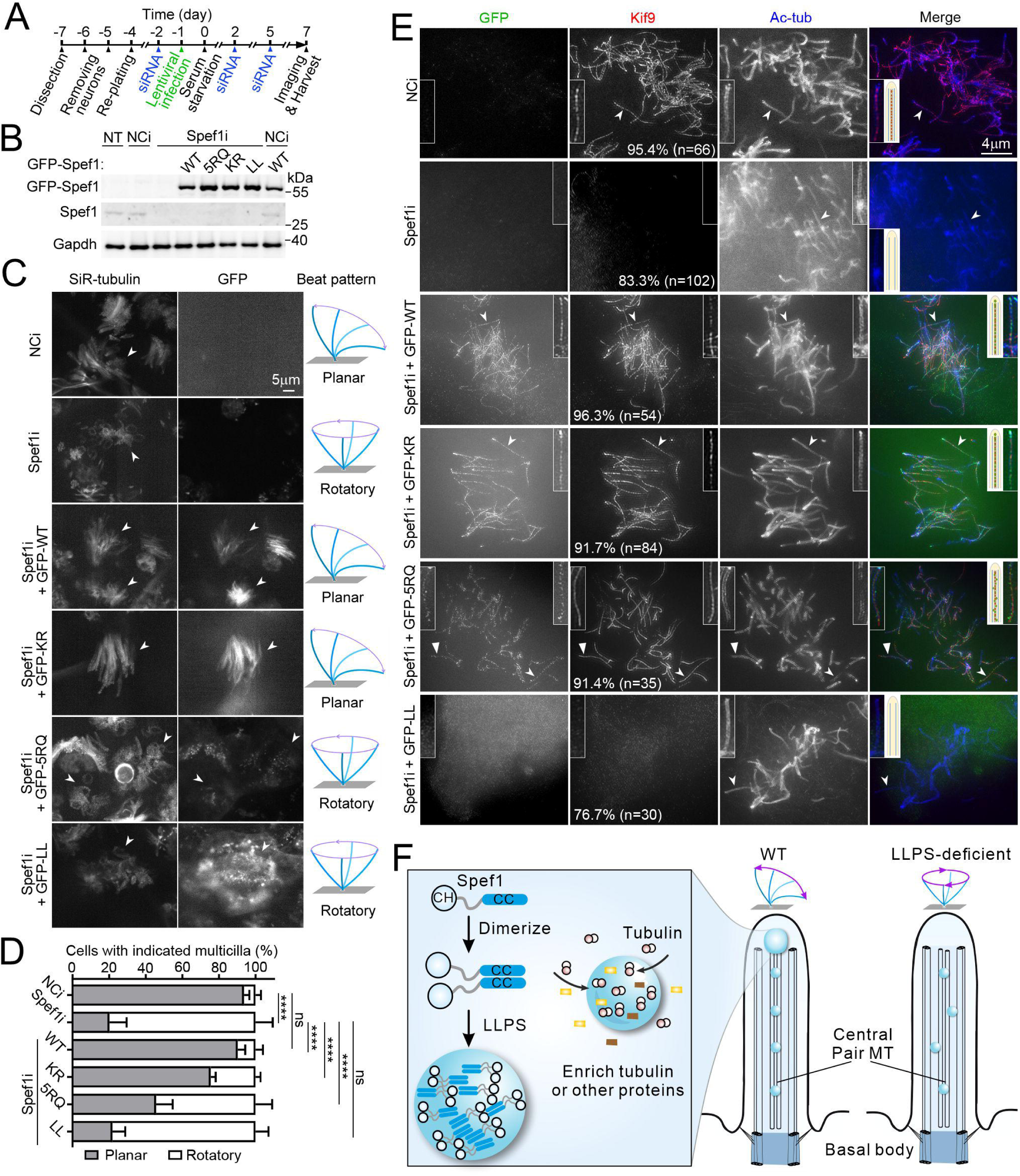
LLPS of Spef1 is critical for its ciliary function. **(A)** Experimental scheme for RNAi and rescue in mEPCs. Glial cells from dissected mouse telencephalon tissues were cultured and induced at day 0 to differentiate into multiciliated mEPCs through serum starvation. The cells were transfected with siRNA and assayed at day 7. GFP-tagged Spef1 or mutants were expressed through lentiviral infections in rescue experiments. **(B)** Expression of RNAi-insensitive Spef1 (WT) and mutants (5RQ, KR and LL) in Spef1-depleted mEPCs. The samples were immunoblotted with anti-Spef1 antibody. Gapdh served as the loading control. NT, non-transfected control; NCi, control siRNA; Spef1i, Spef1 siRNA. **(C-D)** Rescue effects of WT and mutants on ciliary beat pattern. Live imaging was performed to trace motilities of SiR-tubulin-labeled multicilia at 10 ms-intervals in GFP-positive mEPCs. The first 100 frames were overlaid to visualize ciliary beat patterns. Representative overlaid images, together with images of the GFP channel and illustrations for beat patterns, are presented in (C). Quantification results (mean ± SD) (D) were from three independent experiments. At least 50 multiciliated cells were scored in each experiment and condition. One-way ANOVA and Tukey test: ns, no significance (p>0.05); **p<0.01; ***p<0.001; ****p<0.0001. **(E)** Rescue effects of WT and mutants on the CP. mEPCs were immunostained to visualize GFP, acetylated tubulin (Ac-tub; axonemal marker) and Kif9 (CP marker) and subjected to super-resolution microscopy. Representative micrographs are shown. Typical cilia (arrows) were magnified by 2-fold to show details. Illustrations are provided to aid comprehension. **(F)** A schematic working model for Spef1 functions in the CP. The basic functional unit of Spef1 is a dimer. Spef1 dimers phase separate into condensates to enrich tubulins for the continuous growth and extension of CP-MTs. The condensates also bundle, stabilize and maintain CP-MTs to ensure proper functions of the CP apparatus. Refer to Discussion for details.

We next examined detailed ciliary localizations of these mutants and their abilities to rescue the CP through super-resolution immunofluorescent microscopy. We used the kinesin-9 family member Kif9, an evolutionarily-conserved CP-associated protein ^32, 33^, as a CP marker. In multicilia of control (e.g., NCi-treated) mEPCs, Kif9 localized to the CP as a string of puncta in the central lumen of axonemes indicated by acetylated tubulin (Ac-tub) (**Figure 5E**) ^32^. Consistent with the essential role of Spef1 in the CP-MT formation ^10^, the depletion of Spef1 abolished the CP-localization of Kif9, which was rescued by GFP-WT (**Figure 5E**). GFP-KR and GFP-5RQ also rescued the CP-localization of Kif9 (**Figure 5E**). Nevertheless, they displayed different ciliary localizations. Similar to GFP-WT, GFP-KR emerged as puncta co-aligned with the string of Kif9 puncta in the axonemal central lumen.

In contrast, although GFP-5RQ also localized in the central lumen as puncta, many of the puncta deviated away from the central line defined by the Kif9 puncta (**Figure 5E**). In sharp contrast, GFP-LL neither localized in the cilia nor rescued the CP-localization of Kif9 (**Figure 5E**). These results suggest that the KR and 5RQ mutants still support the CP formation, with the 5RQ mutant displaying defected associations with the CP, whereas the LL mutant is non-functional.

Taken together, we conclude that the functional unit of Spef1 is a dimer. Disruption of the dimerization abolishes its ability to undergo LLPS and support the CP formation. Attenuating the LLPS of Spef1 by charge-changing point mutations (as in the 5RQ mutant) while retaining its dimerization does not impair its activity to support the CP formation. The mutant, however, is unable to stably associate with the CP, leading to malfunctions of the CP apparatus.

## Discussion

The axoneme of motile cilia is a MT-based structure featured with a pair of CP-MTs for dictating ciliary beating. Spef1 is a MT-bundling protein essential for the proper assembly of CP-MTs and the central apparatus in the axoneme to control ciliary motility ^10, 24^, although the underlying mechanism remains elusive. In this work, we biochemically characterize Spef1 and reveal its unexpected LLPS capacity *in vitro* and *in cellulo* (**Figure 1**). The Spef1-LLPS condensates can recruit and enrich tubulins to promote MT growth (**Figure 2**). Together with the bundling activity of Spef1, MTs generated from Spef1-LLPS condensates are thick bundled MTs and tend to organize a complex MT-network, suggesting the MT-organization capacity of Spef1. Although Spef1 contains both the IDR and CC domain that are potential LLPS-inducing factors, the CC domain alone is sufficient to drive the LLPS of Spef1 (**Figure 3**). The structure of Spef1-CC reveals a unique charge distribution pattern on the surface (**Figure 4**), in addition to the canonical dimerization capacity, together contributing to the multivalent interactions for initiating LLPS. Based on the LLPS-disrupting mutations, the LLPS capacity of Spef1 is essential for Spef1-mediated ciliary function (**Figure 5**). Thus, the previously unknown LLPS capacity of Spef1 induces the formation of biomolecular condensates with the intrinsic ability to dictate MT growth and organization, which provides a mechanistic explanation for Spef1-mediated control of the CP-MT organization and function in the axoneme of motile cilia.

Spef1-CC is the LLPS-determining site in Spef1 and possesses the dimerization capacity and a special charge distribution pattern both essential for the LLPS of Spef1 (**Figure 3** and **4**). In addition to LLPS, the dimerization capacity of Spef1-CC would also contribute to the MT-bundling activity of Spef1 ^10, 23^, and Spef1-CC thus most likely exhibit the dual functions for LLPS and MT-bundling. At the current stage, we hardly distinguish the versatile functions of Spef1-CC and prefer the scenario in which the LLPS and MT-bundling capacities of Spef1-CC synergistically control MT assembly and organization, as demonstrated by the thick bundled MT-asters generated from Sef1-LLPS condensates (**Figure 2**). Although the dimerization capacity and charge distribution pattern of Spef1-CC are two LLPS-determinants, the CC dimer formation seems to be a prerequisite and the charge distribution pattern is more likely to be an auxiliary factor due to the more severe impact of the CC-disrupting mutation on LLPS (**Figure 4**). Based on the structural analysis of Spef1-CC, the charge distribution pattern on the surface needs the pre-formed CC dimer (**Figure 4**), and the disruption of the CC dimer not only impair the dimerization capacity but also change the charge distribution pattern, which explains the prerequisite role of the CC dimer formation for LLPS.

In accordance with the prominent role of Spef1-CC for LLPS, the LLPS-disrupting mutations in Spef1-CC severely impair Spef1-mediated ciliary functions (**Figure 5A-E**). More intriguingly, the two types of mutations (CC-disrupting and charge-changing) in Spef1-CC also show distinct impacts on Spef1 functions. The CC-disrupting (L208A/L212A) mutant is completely inactive (**Figure 5C-E**), indicating the importance of Spef1 dimerization. Previous publications indicate that the dimerization enables Spef1 to stabilize MTs ^10^. As the dimerization is essential for the LLPS of Spef1 (**Figure 4**), we propose that the LLPS also contributes to the CP formation by facilitating MT polymerization (**Figure 5F**). Interestingly, although the charge-changing 5RQ mutant displays markedly reduced LLPS *in vitro* (**Figure 4G**), it still supports the CP formation and forms puncta in the central lumen (**Figure 5E**), suggesting the preservation of some *in vivo* LLPS ability. Nevertheless, the puncta fail to tightly associate with the CP, and the CP apparatus is not fully functional because a large portion of the cilia display a rotatory beat pattern (**Figure 5C-E**), suggesting that the proper localization of the Spef1 puncta to the CP seems to be also essential for Spef1-mediated ciliary functions in motile cilia.

LLPS is a process of demixing components from solution to form biomolecular condensates for controlling a variety of cellular activities ^30, 34^. A number of MT-binding proteins are capable of undergoing LLPS to regulate MT-related processes such as nucleation, dynamics and organization ^35^. MT-plus-end-binding EB proteins can form condensed droplets through LLPS that recruit MT-binding regulators to constitute a machinery for guiding MT dynamics ^36, 37^, while MT-minus-end-binding Camsap proteins are able to co-condense with tubulins via LLPS to organize a nucleation center for dictating non-centrosomal MT growth ^38^. Moreover, the LLPS condensates of some MT-associated proteins can even act as a prominent nucleation and branching center along the MT lattice to initiate its branching for a network organization ^39, 40^. We here identify Spef1 as a new member of MT-associated proteins with the LLPS capacity to form biomolecular condensates. Similar to Camsap proteins, the Spef1-LLPS condensates can enrich tubulins and generate thick aster-like MT-bundle structures (**Figure 2**), suggesting that these condensates may also function as a nucleation center to initiate the growth and organization of non-centrosomal MTs. Since Spef1 and Camsap proteins are both essential for proper non-centrosomal CP-MT formation ^9, 10^, these two types of LLPS-capable proteins are most likely to work together to organize CP-MTs in the axoneme of motile cilia. Thus, LLPS may be a common process for generating and organizing non-centrosomal CP-MTs.

We thus propose a working model for Spef1-mediated regulation of CP-MT formation in the axoneme based on available biochemical and cellular data. In nascent cilia, Spef1 is predominantly enriched at the tip of cilia to form biomolecular condensates through LLPS (**Figure 5F**). The Spef1-LLPS condensates would then enrich tubulins for the continuous growth and extension of CP-MTs. Along with the extension of CP-MTs, Spef1 condensates would further bundle and stabilize CP-MTs and prevent the unfavorable disassembly (**Figure 5F**). In mature cilia, Spef1 condensates, present as puncta along the CP, would also work as MT nucleation centers and tubulin sources to repair CP-MTs damaged during repeated ciliary beating (**Figure 5F**). Moreover, these Spef1 puncta tightly associated with and distributed along the CP would likely function as buttresses to stabilize CP-MTs to accommodate the external force and coordinate planar ciliary beating. In this context, the abnormal ciliary beat in mEPCs expressing the 5RQ mutant (**Figure 5C-D**) can be attributed to repair defects of damaged CP-MTs or the partial impairment of the buttress-like function. It will be interesting to verify the proposed mechanisms in future investigations.

## Materials and Methods

### Protein expression and purification

DNA sequences encoding mouse Spef1 or various Spef1 fragments were each cloned into a modified pET32a vector with an N-terminal GFP-His_6_-tag or GB1-His_6_-tag for the expression of GFP-tagged proteins or untagged proteins, respectively. Recombinant proteins were expressed in *Escherichia coli* BL21(DE3) host cells at 16°C. Various Spef1 fragments were purified by Ni^2+^-Sepharose 6 Fast Flow (GE healthcare) affinity chromatography followed by size-exclusion chromatography (Superdex-200 26/60, GE healthcare) with the buffer containing 50 mM Tris, pH 8.0, 500 mM NaCl, 1 mM EDTA, 1 mM DTT. For untagged proteins, another step of size-exclusion chromatography (Superdex-200 26/60, GE healthcare) was performed after the cleavage of the tag.

### Cell culture and transfection

The full-length wild-type Spef1 and various mutants were each cloned into a pEGFP-C3 vector. The coiled-coil domain of Spef1 was cloned into a pMKO-N1 vector. HeLa, HEK293 and COS-7 cells were maintained in Dulbecco’s modified Eagle medium (Gibco) with 10% (v/v) foetal bovine serum (FBS, Hyclone) and 100 U/ml penicillin (Gibco) plus 100μg/ml streptomycin (Gibco). Plasmids were purified with endotoxin-free purification kits (CoWin Biosciences) and transfected into cells with Lipofectamine 3000 (Invitrogen) according to the user manual.

### FRAP experiment

FRAP experiments were performed on an inverted laser scanning confocal microscope (Zeiss, LSM980 AxioObserver.Z1/7). For living cell imaging, cell culture dishes were placed in an incubation chamber maintained at 37°C with 5% (v/v) CO_2_. For *in vitro* FRAP experiment, samples were also placed in the same chamber maintained at set temperature. Images were acquired with a Plan-Apochromat 63×/1.40 NA oil immersion objective under the control of the Zeiss Zen software. FRAP experiment images were acquired for two time points before bleaching and then by bleaching with 100% intensity of the 488-nm or 555-nm laser and imaging every 2 s for 15 min or 20 min. The fluorescence intensity of the bleached area was normalized to the intensity of the same area before bleaching, and the recovery curves were corrected with the region without photobleaching in the same frames. Intensity traces were analyzed using Imaris9.1 software (Oxford Instrument) or ImageJ (NIH).

### *In vitro* phase separation assay

Phase separation was induced by diluting the indicated amount of Spef1 in the low salt buffer for phase separation reactions. Droplet formation was observed by adding a drop of the reaction solution onto a glass-bottom culture dish (NEST Biotechnology) and then imaging the droplet on a Zeiss LSM 980 confocal microscope with a Plan-Apochromat 63×/1.4 NA oil immersion lens by fluorescence and differential interference contrast imaging.

### Tubulin purification and labeling

Crude tubulins were obtained from porcine brain through double cycles of polymerization and depolymerization, as previously described ^41^. Tubulins were further purified using a TOG-based affinity column ^42^. Tubulins were labeled with TAMRA (Thermo Fisher Scientific) using NHS esters, following standard protocols ^43^.

### Tubulin polymerization assay

Tubulin polymerization assays were performed by using the tubulin polymerization assay kit (Cytoskeleton, Inc.). Briefly, 30 μM of tubulins and the indicated concentrations of Spef1 were mixed on ice in PEM buffer (80 mM PIPES, pH 6.9, 1 mM EGTA, 1 mM GTP, and 1 mM MgCl_2_) to a total volume of 100 μl. Then the mixture was added to a 96-well plate pre-warmed at 37°C. Absorbance at 340 nm was immediately measured with a Varioskan Flash spectrophotometer (Thermo Fischer Scientific) every 30 s for 61 min.

### Crystallization, data collection and structural determination

Purified Spef1-CC was concentrated to ∼10 mg/ml in the buffer containing 50 mM Tris-HCl pH 8.0, 500 mM NaCl, 1 mM EDTA, and 1 mM DTT for crystallization. Crystals of Spef1-CC were grown in 0.1 M BICINE, pH 8.5, 3% (v/v) Dextran sulfate sodium salt and 15% (w/v) PEG 20000 using the vapor diffusion method (sitting drop) at 16°C. Before being flash-frozen in liquid nitrogen, crystals were cryo-protected in their mother liquor supplemented with 16% (v/v) ethylene glycol. Diffraction data were collected at the beamline BL19U at the Shanghai Synchrotron Radiation Facility (SSRF) using with a wavelength of 0.979 Å at 100K ^44^, and were processed and scaled using HKL2000 ^45^. The structure was determined by the molecular replacement method with idealized coiled-coil structures as searching models using AMPLE ^46^. The sidechains were manually modeled into the structure according to the *2Fo-Fc* and *Fo-Fc* electron density maps using COOT ^47^. The structure was further fitted and refined with PHENIX ^48^. The structure figures were prepared with the program ChimeraX ^49^. The statistics for data collection and structural refinement were summarized in **Table S1**.

### Size exclusion chromatography coupled with multi-angle light scattering (SEC-MALS)

Protein samples (∼1 mg/ml in 50 mM Tris-HCl pH 8.0, 500 mM NaCl, 1 mM EDTA, and 1 mM DTT) were analyzed with static light scattering by injection of them into an Agilent FPLC system with a WTC SEC column (Wyatt Technology). The chromatography system was coupled with an 18-angle light-scattering detector (DAWN HELEOS II, Wyatt Technology) and differential refractive index detector (Optilab rEx, Wyatt Technology). Masses (molecular weights) were calculated with ASTRA (Wyatt Technology). Bovine serum albumin (Sigma) was used as the calibration standard.

### Electron microscopy

Samples, including tubulins, Spef1 or Spef1 mutants and the mixture of them, were diluted with PEM buffer (80 mM PIPES pH 6.9, 1 mM MgCl_2_, 1 mM EGTA, and 1 mM GTP) and incubated on ice for 10 min and then placed in 37°C water bath for 30 min before blotting.

Four microliters of co-polymerized samples were incubated on a glow-discharged copper grid (200 mesh, Beijing Zhongjingkeyi Technology Co., Ltd) for 1 min and then washed by an aliquot of deionized water and 3% w/v uranyl acetate, followed by staining with an aliquot of 3% w/v uranyl acetate for another 42 s and dried in air. The specimens were observed using a Tecnai G2 Spirit transmission electron microscope (FEI Company, 100kV).

### *In vitro* MT-bundling assay

MTs were polymerized *in vitro*. Briefly, 5 μl of porcine tubulin stock (5mg/ml, the ratio of tubulin to TAMRA-labeled tubulin is 19:1) in the reaction buffer (80 mM PIPES, pH 6.9, 1 mM GTP, 2 mM MgCl_2_, 0.5 mM EGTA, 40% glycerol) was incubated at 37°C overnight, then diluted with 195 μl of pre-warmed (25°C) stabilization buffer (80 mM PIPES, pH 6.9, 20 μM Taxol, 2 mM MgCl_2_, 0.5 mM EGTA). To assay for MT-bundling ability, 2 μl of the Taxol-stabilized MTs was mixed with 6 μl purified proteins of varying concentrations. After incubation at 37°C for 10 min, 3 μl of each mixture were gently squashed under an 18-mm circular coverslip. Microscopic examination and imaging were performed using Olympus FV1200 microscope with a 100×/1.35 oil immersion objective.

### mEPC cell culture, transfection and infection

Multiciliated mEPCs were obtained and cultured as described ^21^. The telencephalon of each P0 mouse was dissected in cold dissection solution (161 mM NaCl, 5 mM KCl, 1 mM MgSO_4_, 3.7 mM CaCl_2_, 5 mM HEPES, and 5.5 mM Glucose, pH 7.4) and digested with 1 ml of the dissection solution (10 U/ml papain (Worthington), 0.2 mg/ml L-Cysteine, 0.5 mM EDTA, 1 mM CaCl_2_, 1.5 mM NaOH) for 30 min at 37°C. Then the dissection solution was replaced with culture medium (Dulbecco’s Modified Eagle medium (DMEM, Gibco) supplemented with 10% fetal bovine serum (Gibco), and 0.1% Primocin (Invivogen)). After gentle pipetting and centrifugation at 1400 rpm for 5 min at room temperature, the cells were resuspended in culture medium and plated on a fibronectin (Millipore)-coated flask (Corning) and cultured in the culture medium at 37°C in 5% (v/v) CO_2_. Neurons were removed by flapping the flask at 24 h and 48 h. The remaining glial cells were cultured to approximately 80% confluency and transferred into fibronectin-coated 29-mm glass-bottom dishes (Cellvis), followed by serum starvation with a serum-free medium to induce multiciliation.

Validated Spef1 siRNA (5’-CAACUUAAGAACGUGCGCAtt-3’) or a control siRNA (5’-UUCUCCGAACGUGUCACGUtt-3’) ^10^ was transfected into mEPCs via Lipofectamine RNAiMAX (Life Technologies). For each 29-mm glass-bottom dish, 3 μl RNAiMAX and 2 pmol siRNA were diluted in 125 μl opti-MEM (Thermo Fisher). After 15 min, the mixed solution was dropped into the dish containing 750 μl fresh medium. The medium was refreshed after 24 h. For rescue experiments, RNAi-insensitive Spef1 constructs were constructed by mutating the siRNA-targeting region (ACTTAAGAACGTGCGC → GCTCAAAAATGTACGT) by PCR using Mutagenesis Kit (Toyobo). Lentiviral particles were packaged by transfecting lentiviral expression plasmids and packaging plasmids, pCMVdr8.9 and pMD2.VSVG, into HEK293T cells. After 48 h, the culture medium containing lentiviral particles was centrifuged at 300×g for 15 min to discard cell debris. The supernatant was used to infect mEPCs to express GFP-Spef1 or mutants.

### Live imaging

For live imaging, multicilia of mEPCs cultured to day 7 were labeled with SiR-tubulin (100 nM final concentration; Spirochrome) in serum-free medium for 1 h at 37°C. After a snapshot of z-slices to locate GFP-positive cells, ciliary motilities at an appropriate z-plane were recorded at 10 ms-intervals for 200 frames by using an Olympus SpinSR microscope with a 60×/1.42 NA oil immersion objective. Image sequences were processed using Fiji (NIH) ^50^. The first 100 frames were overlaid to generate ciliary beat tracks for quantifications of ciliary beat patterns. Cilia with linear beat tracks were considered to beat in a planar pattern, whereas those with circular beat tracks were scored as a rotatory pattern. mEPCs with multicilia of mixed beat patterns were scored according to behaviors of the majority of the cilia.

### Immunofluorescent staining and imaging

mEPCs cultured to day 7 were washed with PBS and pre-permeabilized with 0.5% Triton X-100 in PBS for 30 s to remove soluble proteins, followed by fixation with 4% paraformaldehyde in PBS for 15 min at room temperature. After permeabilization with 0.5% Triton X-100 in PBS for 15 min, the samples were blocked with 4% BSA in PBS for 1 h and incubated with primary antibodies overnight and secondary antibodies for 1 h. Washes were performed with 4% BSA in PBS for three times after each antibody incubation. After the last wash with PBS, the samples were mounted with ProLong Diamond anti-fade mounting medium (Invitrogen). Primary antibodies were mouse anti-acetylated tubulin (Sigma-Aldrich), rabbit anti-Kif9 ^32^, and chicken anti-GFP (Invitrogen). Secondary antibodies were goat anti-mouse pacific blue (Invitrogen), goat anti-IgY Alexa Fluor 488 (Invitrogen), and goat anti-rabbit Alexa Fluor 647 (Invitrogen). Images were acquired by using a High Sensitivity Structured Illumination Microscope (HiS-SIM, CSR Biotech) with a Plan Apo 100×/1.5 NA oil-immersion objective lens (Olympus).

### Western blot

mEPCs were lysed with 2× SDS PAGE loading buffer and boiled at 100°C for 10 min after finishing live imaging. After SDS-PAGE and membrane transfer, endogenous and exogenous Spef1 was detected using an anti-Spef1 antibody ^10^, whereas Gapdh was detected using an anti-Gapdh antibody (Proteintech).

### Accession numbers

The atomic coordinate of the coiled-coil domain of Spef1 has been deposited in the Protein Data Bank with the accession code 9IJX.

## Acknowledgements

We thank the Shanghai Synchrotron Radiation Facility for the beam time and Dr. Xiumin Yan for the helpful discussion on this project. This work was supported by grants from the CAS Project for Young Scientists in Basic Research (YSBR-104 and YSBR-075), the National Natural Science Foundation of China (32371273), the Chinese National Programs for Brain Science and Brain-like Intelligence Technology (2022ZD0205800), the Beijing Natural Science Foundation (5242021), and the Youth Innovation Promotion Association of CAS.

## References

1. Satir, P. & Christensen, S.T. Overview of structure and function of mammalian cilia. Annual review of physiology 69, 377–400 (2007).

2. Ishikawa, T. Axoneme Structure from Motile Cilia. Cold Spring Harbor perspectives in biology 9 (2017).

3. Ginger, M.L., Portman, N. & McKean, P.G. Swimming with protists: perception, motility and flagellum assembly. Nature reviews. Microbiology 6, 838–850 (2008).

4. Brooks, E.R. & Wallingford, J.B. Multiciliated cells. Current biology: CB 24, R973–982 (2014).

5. Satir, P., Heuser, T. & Sale, W.S. A Structural Basis for How Motile Cilia Beat. Bioscience 64, 1073–1083 (2014).

6. Loreng, T.D. & Smith, E.F. The Central Apparatus of Cilia and Eukaryotic Flagella. Cold Spring Harbor perspectives in biology 9 (2017).

7. Oda, T., Yanagisawa, H., Yagi, T. & Kikkawa, M. Mechanosignaling between central apparatus and radial spokes controls axonemal dynein activity. The Journal of cell biology 204, 807–819 (2014).

8. Zhu, X., Liu, Y. & Yang, P. Radial Spokes-A Snapshot of the Motility Regulation, Assembly, and Evolution of Cilia and Flagella. Cold Spring Harbor perspectives in biology 9 (2017).

9. Liu, H. et al. Wdr47, Camsaps, and Katanin cooperate to generate ciliary central microtubules. Nature communications 12, 5796 (2021).

10. Zheng, J. et al. Microtubule-bundling protein Spef1 enables mammalian ciliary central apparatus formation. Journal of molecular cell biology 11, 67–77 (2019).

11. Fliegauf, M., Benzing, T. & Omran, H. When cilia go bad: cilia defects and ciliopathies. Nature reviews. Molecular cell biology 8, 880–893 (2007).

12. Praveen, K., Davis, E.E. & Katsanis, N. Unique among ciliopathies: primary ciliary dyskinesia, a motile cilia disorder. F1000prime reports 7, 36 (2015).

13. Lechtreck, K.F., Gould, T.J. & Witman, G.B. Flagellar central pair assembly in Chlamydomonas reinhardtii. Cilia 2, 15 (2013).

14. Dymek, E.E., Lefebvre, P.A. & Smith, E.F. PF15p is the chlamydomonas homologue of the Katanin p80 subunit and is required for assembly of flagellar central microtubules. Eukaryotic cell 3, 870–879 (2004).

15. Dymek, E.E. & Smith, E.F. PF19 encodes the p60 catalytic subunit of katanin and is required for assembly of the flagellar central apparatus in Chlamydomonas. Journal of cell science 125, 3357–3366 (2012).

16. Hao, L. et al. Intraflagellar transport delivers tubulin isotypes to sensory cilium middle and distal segments. Nature cell biology 13, 790–798 (2011).

17. Robinson, A.M. et al. CAMSAP3 facilitates basal body polarity and the formation of the central pair of microtubules in motile cilia. Proceedings of the National Academy of Sciences of the United States of America 117, 13571–13579 (2020).

18. Akhmanova, A. & Steinmetz, M.O. Control of microtubule organization and dynamics: two ends in the limelight. Nature reviews. Molecular cell biology 16, 711–726 (2015).

19. Hendershott, M.C. & Vale, R.D. Regulation of microtubule minus-end dynamics by CAMSAPs and Patronin. Proceedings of the National Academy of Sciences of the United States of America 111, 5860–5865 (2014).

20. Jiang, K. et al. Microtubule minus-end stabilization by polymerization-driven CAMSAP deposition. Developmental cell 28, 295–309 (2014).

21. Ren, J. et al. Intertwined Wdr47-NTD dimer recognizes a basic-helical motif in Camsap proteins for proper central-pair microtubule formation. Cell reports 41, 111589 (2022).

22. Chan, S.W., Fowler, K.J., Choo, K.H.A. & Kalitsis, P. Spef1, a conserved novel testis protein found in mouse sperm flagella. Gene 353, 189–199 (2005).

23. Dougherty, G.W. et al. CLAMP, a novel microtubule-associated protein with EB-type calponin homology. Cell motility and the cytoskeleton 62, 141–156 (2005).

24. Tapia, R. & Hecht, G.A. Spef1/CLAMP binds microtubules and actin-based structures and regulates cell migration and epithelia cell polarity. Annals of the New York Academy of Sciences 1515, 97–104 (2022).

25. Werner, M.E. et al. Radial intercalation is regulated by the Par complex and the microtubule-stabilizing protein CLAMP/Spef1. Journal of Cell Biology 206, 367–376 (2014).

26. Kim, S.K. et al. CLAMP/Spef1 regulates planar cell polarity signaling and asymmetric microtubule accumulation in the Xenopus ciliated epithelia. The Journal of cell biology 217, 1633–1641 (2018).

27. Gray, R.S. et al. The planar cell polarity effector Fuz is essential for targeted membrane trafficking, ciliogenesis and mouse embryonic development. Nature cell biology 11, 1225–1232 (2009).

28. Wieczorek, M., Bechstedt, S., Chaaban, S. & Brouhard, G.J. Microtubule-associated proteins control the kinetics of microtubule nucleation. Nature cell biology 17, 907–916 (2015).

29. Lukinavicius, G. et al. Fluorogenic probes for live-cell imaging of the cytoskeleton. Nature methods 11, 731–733 (2014).

30. Banani, S.F., Lee, H.O., Hyman, A.A. & Rosen, M.K. Biomolecular condensates: organizers of cellular biochemistry. Nature reviews. Molecular cell biology 18, 285–298 (2017).

31. Dignon, G.L., Best, R.B. & Mittal, J. Biomolecular Phase Separation: From Molecular Driving Forces to Macroscopic Properties. Annual review of physical chemistry 71, 53–75 (2020).

32. Fang, C. et al. Distinct roles of Kif6 and Kif9 in mammalian ciliary trafficking and motility. The Journal of cell biology 223 (2024).

33. Han, L. et al. Cryo-EM structure of an active central apparatus. Nat Struct Mol Biol 29, 472–482 (2022).

34. Hyman, A.A., Weber, C.A. & Julicher, F. Liquid-liquid phase separation in biology. Annual review of cell and developmental biology 30, 39–58 (2014).

35. Volkov, V.A. & Akhmanova, A. Phase separation on microtubules: from droplet formation to cellular function? Trends in cell biology (2023).

36. Song, X. et al. Phase separation of EB1 guides microtubule plus-end dynamics. Nature cell biology 25, 79–91 (2023).

37. Meier, S.M. et al. Multivalency ensures persistence of a +TIP body at specialized microtubule ends. Nature cell biology 25, 56–67 (2023).

38. Imasaki, T. et al. CAMSAP2 organizes a gamma-tubulin-independent microtubule nucleation centre through phase separation. eLife 11 (2022).

39. Jijumon, A.S. et al. Lysate-based pipeline to characterize microtubule-associated proteins uncovers unique microtubule behaviours. Nature cell biology 24, 253–267 (2022).

40. King, M.R. & Petry, S. Phase separation of TPX2 enhances and spatially coordinates microtubule nucleation. Nature communications 11, 270 (2020).

41. Gell, C. et al. Purification of tubulin from porcine brain. Methods in molecular biology 777, 15–28 (2011).

42. Widlund, P.O. et al. One-step purification of assembly-competent tubulin from diverse eukaryotic sources. Molecular biology of the cell 23, 4393–4401 (2012).

43. Hyman, A. et al. Preparation of modified tubulins. Methods in enzymology 196, 478–485 (1991).

44. Wang, Q.S. et al. Upgrade of macromolecular crystallography beamline BL17U1 at SSRF. Nucl Sci Tech 29 (2018).

45. Otwinowski, Z. & Minor, W. Processing of X-ray diffraction data collected in oscillation mode. Methods in enzymology 276, 307–326 (1997).

46. Bibby, J., Keegan, R.M., Mayans, O., Winn, M.D. & Rigden, D.J. AMPLE: a cluster-and-truncate approach to solve the crystal structures of small proteins using rapidly computed ab initio models. Acta crystallographica. Section D, Biological crystallography 68, 1622–1631 (2012).

47. Emsley, P. & Cowtan, K. Coot: model-building tools for molecular graphics. Acta crystallographica. Section D, Biological crystallography 60, 2126–2132 (2004).

48. Adams, P.D. et al. PHENIX: a comprehensive Python-based system for macromolecular structure solution. *Acta crystallographica. Section D*, Biological crystallography 66, 213–221 (2010).

49. Pettersen, E.F. et al. UCSF ChimeraX: Structure visualization for researchers, educators, and developers. Protein science: a publication of the Protein Society 30, 70–82 (2021).

50. Schindelin, J. et al. Fiji: an open-source platform for biological-image analysis. Nature methods 9, 676–682 (2012).

