## Supplemental figures and table for "Coiled-coil-mediated phase separation of Spef1 for non-centrosomal microtubule organization and function"

**Supplementary figures and legends**

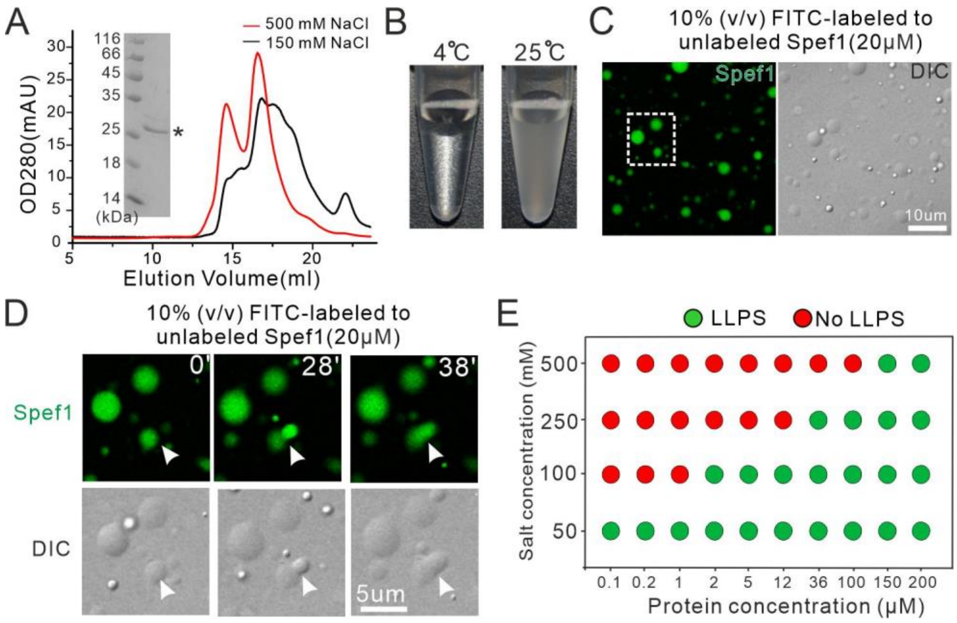


**Figure S1: Temperature- and salt concentration-dependent LLPS of Spef1. (A)** Salt concentration-dependent behaviors of Spef1 characterized by the gel-filtration analysis. The inset shows the SDS-PAGE analysis of Spef1 purified from *E. coli*. **(B)** Temperature dependence of the phase separation of Spef1. Spef1 exhibits phase separation behavior under elevated temperature conditions. **(C)** Representative fluorescence and DIC images of the phase separation of Spef1. Prior to the experiment, the FITC-labeled protein was mixed with the unlabeled protein at a molar ratio of 1:9 (labeled: unlabeled). **(D)** Fusion of Spef1 droplets with the increase of time. **(E)** LLPS diagram showing the phase separation of Spef1 at different protein and salt concentrations.

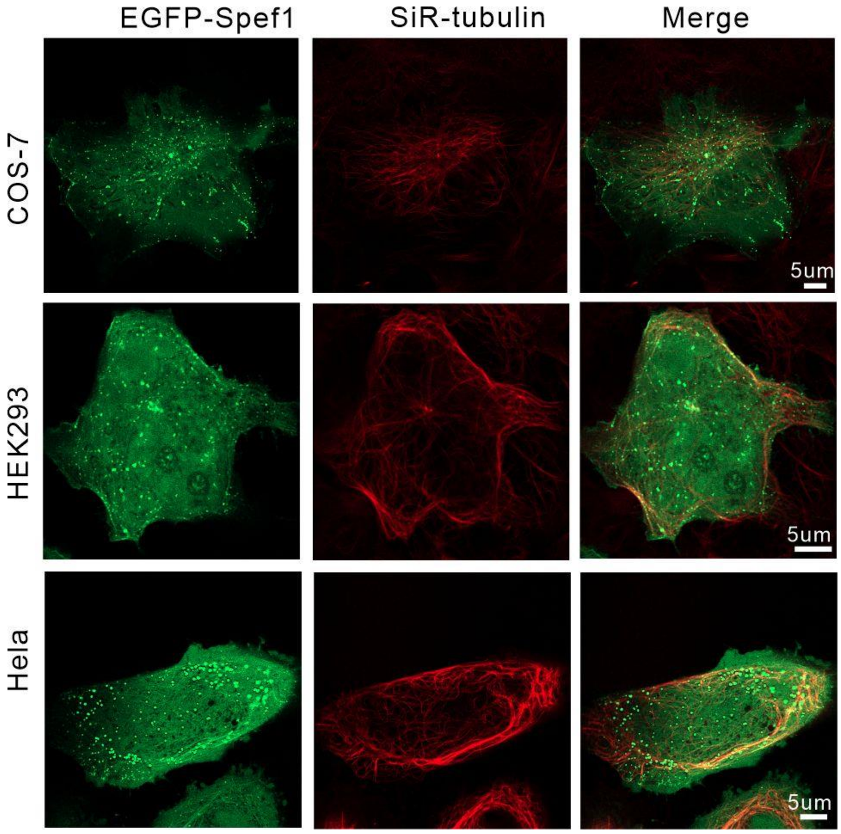


**Figure S2: Exogenous Spef1 expressed in different cell lines**. Representative fluorescence images of COS-7, HEK293 and Hela cells transfected with pEGFP-Spef1 to express Spef1 tagged with EGFP. Cellular microtubules were labeled with SiR-tubulin. Scale bar: 5 μm.

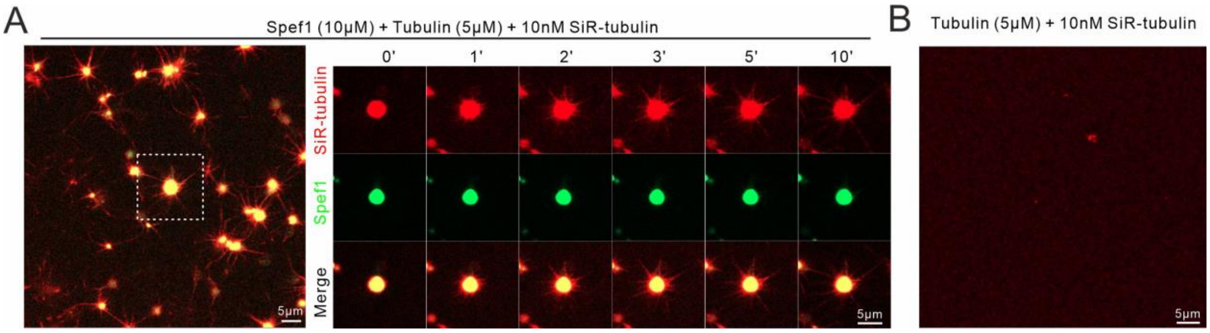


**Figure S3: Spef1 condensates enrich unlabeled tubulins to promote MT assembly. (A)** Left: Representative fluorescence microscopy of EGFP-Spef1 (green) and unlabeled tubulins taken 10 min after starting the reaction with the addition of the microtubule dye SiR-tubulin. Right: Time-lapse fluorescent images of the mixture of EGFP-Spef1 and unlabeled tubulins. **(B)** Representative fluorescence microscopy of unlabeled tubulins taken 10 min after starting the reaction. Scale bar: 5 μm.

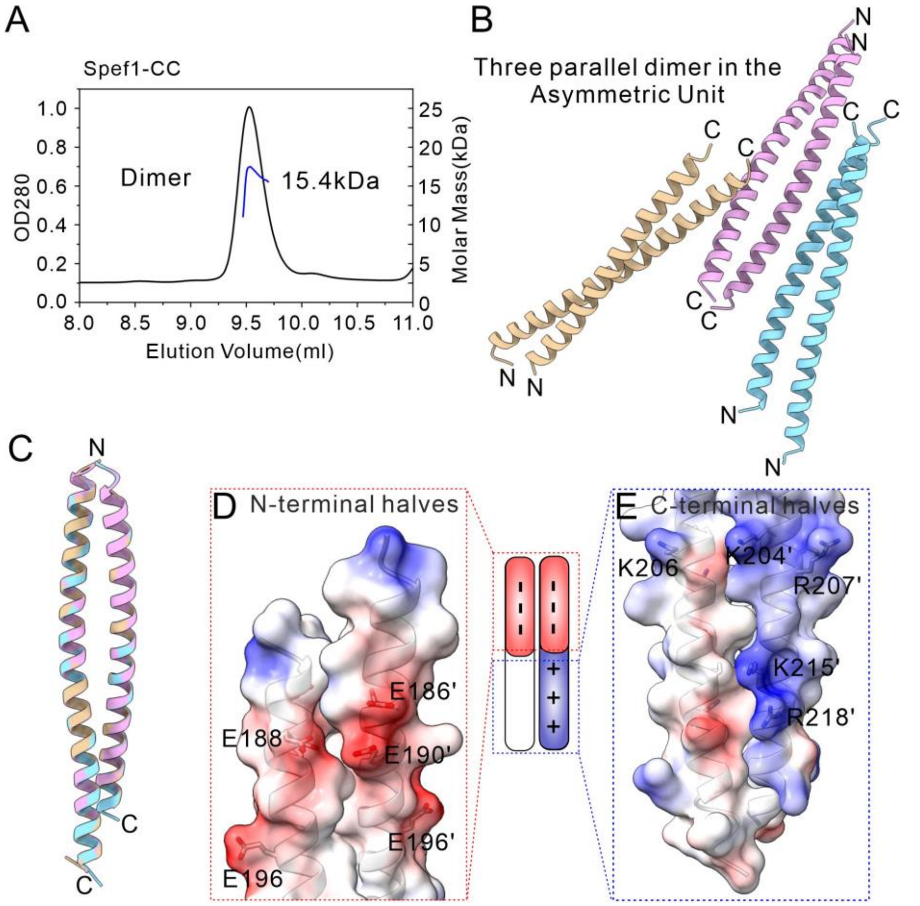


**Figure S4: Structure of the Spef1-CC dimer. (A)** Biochemical characterization of the oligomeric states of Spef1-CC. **(B)** Overall structure of the Spef1-CC dimer in the asymmetric unit of the crystal packing. Three parallel dimers packs in the asymmetric unit. **(C)** Structural super-imposition of the three dimers in the asymmetric unit. **(D-E)** Electrostatic surface combined with stick model showing the location of the negatively charged residues (E186, E188, E190 and E196) and the positively charged residues (K204, K206, R207, K215 and R218).

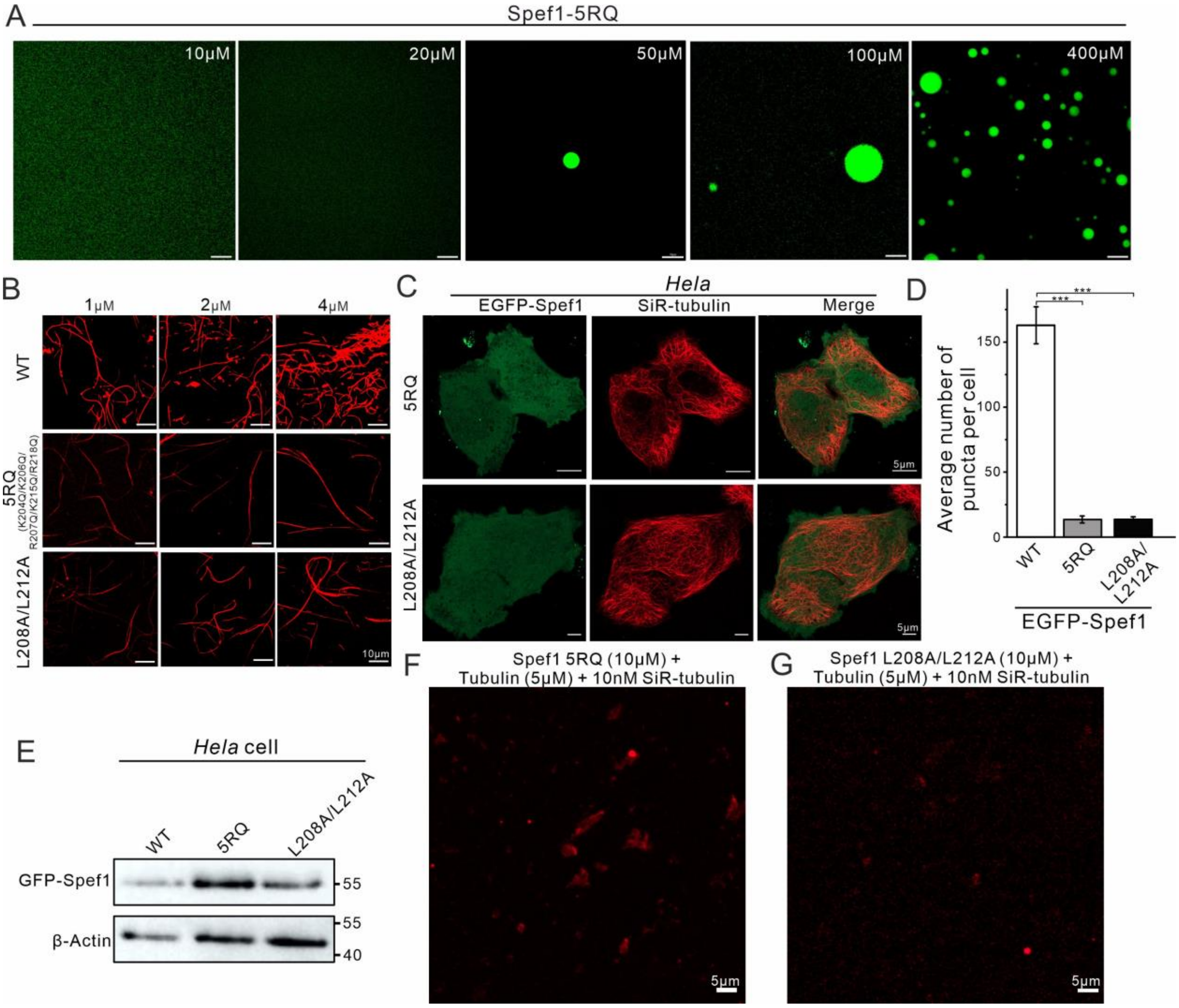


**Figure S5: Characterization of the MT-bundling abilities of the different Spef1 mutants *in vitro* and the LLPS capabilities of them *in cellulo*. (A)** In-vitro phase separation behaviors of GFP-Spef1-5RQ mutant. The concentration of protein used for LLPS induction were labeled in the top-left corner of the panel. Scale bar: 10 μm. **(B)** *In vitro* MT-bundling assay. MTs polymerized *in vitro* were incubated with the indicated concentrations of the wild-type (WT) Spef1 and its mutants at 37°C for 10 min for imaging. **(C)** Confocal images of Hela cells expressing GFP-Spef1-5RQ and GFP-Spef1-L208/L212A. MTs were labeled with SiR-Tubulin. Scale bar: 5 μm. **(D)** Quantification of the average number of puncta per cell. Data represent mean ± SEM; statistical significance was determined using Student's t-test (***$p < 0.001$). **(E)** Stability analysis of EGFP-Spef1 wild-type (WT), 5RQ, and LL mutants by Western blotting. Plasmids encoding EGFP-tagged wild-type or mutant Spef1 were transfected into *HeLa* cells. Thirty-three hours post-transfection, cell lysates were subjected to Western blot analysis using an anti-GFP antibody. β-actin was used as a loading control. **(F-G)** Representative fluorescence microscopy of EGFP-Spef1 with the 5RQ (F) and L208A/L212A mutations (G) and unlabeled tubulins taken 10 min after starting the reaction. Scale bar: 5 μm.

**Table S1. Data collection and structural refinement statistics**

| A. Diffraction data | Spef1 Coiled coil (PDB code: 9IJX) |
|---|---|
| Space group | C2 |
| Wavelength (Å) | 0.979 |
| Cell dimensions | |
| a, b, c (Å) | 60.2, 104.4, 72.3 |
| α, β, γ (º) | 90, 90, 90 |
| Resolution (Å) | 50.0-2.45(2.51-2.45) [a] |
| Unique reflections | 15881(1167) |
| I/σ(I) | 11.4(2.4) |
| Multiplicity | 4.4(3.6) |
| Completeness (%) | 96.6(93.7) |
| B. Refinement | 26.09-2.45(2.60-2.45) |
| Rwork (%) | 19.5 |
| Rfree (%) | 23.4 |
| Mean B factors ($\text{Å}^2$) | 97.0 |
| R.m.s. deviation[b] | |
| Bond length (Å) | 0.012 |
| Bond angles (º) | 1.189 |
| Ramachandran plot (%) | |
| Favored region | 95.8 |
| Allowed region | 4.2 |
| Disallowed region | 0 |

[a]The values in parentheses refer to the highest resolution shell.

[b]Root mean square deviation from ideal values.

**Video S1: Time-lapse movie of MT-aster formation in Spef1 droplets.** Precooled tubulins (5 µM), SiR-tubulin (10 nM), GTP (1 mM), $MgCl_2$ (4 mM), and GFP-Spef1 (10 µM) were mixed and transferred to a chamber incubated at 37 C. Video recording began immediately after setup, with a frame rate of 1 frame per second.
